# PD-L1 glycosylation and its impact on binding to clinical antibodies

**DOI:** 10.1101/2020.07.03.187179

**Authors:** Julius Benicky, Miloslav Sanda, Zuzana Brnakova Kennedy, Oliver C. Grant, Robert J. Woods, Alan Zwart, Radoslav Goldman

**Affiliations:** Department of Oncology, Lombardi Comprehensive Cancer Center, Georgetown University. Washington, D.C. 20057; Department of Biochemistry and Molecular & Cellular Biology, Georgetown University, Washington, DC, 20057; Clinical and Translational Glycoscience Research Center, Georgetown University, Washington, DC, 20057; Department of Biochemistry and Molecular Biology, Complex Carbohydrate Research Center, University of Georgia, Athens, GA 30602

**Keywords:** programmed cell death ligand-1, PD-L1, N-glycosylation, site-specific glycosylation, polyLacNAc, immune checkpoints, therapeutic antibody, durvalumab, binding constant, surface plasmon resonance

## Abstract

Immune checkpoint inhibitors, including PD-L1/PD-1, are key regulators of immune response and promising targets in cancer immunotherapy. N-glycosylation of PD-L1 affects its interaction with PD-1 but little is known about the distribution of glycoforms at its four NXS/T sequons. We optimized LC-MS/MS methods using collision energy modulation for the site-specific resolution of specific glycan motifs. Using these methods, we demonstrate that PD-L1 expressed on the surface of breast cancer cells carries mostly complex glycans with high proportion of polyLacNAc structures at the N219 sequon. PD-L1 from whole cell lysate contained, in addition, large proportion of high mannose glycans at all sites. Contrary to the full-length protein, the secreted form of PD-L1 expressed in breast cancer or HEK293 cells demonstrated minimum N219 occupancy and low contribution of the polyLacNAc structures. Molecular modeling of PD-L1/PD-1 interaction with N-glycans suggests that glycans at the N219 site of PD-L1 and N74 and N116 of PD-1 are involved in glycan-glycan interactions, but the impact of this potential interaction on the protein function remains at this point unknown. In addition, the interaction of PD-L1 with clinical antibodies is also affected by glycosylation. In conclusion, our study demonstrates that PD-L1 expressed in the MDA-MB-231 breast cancer cells carries polyLacNAc glycans mostly at the N219 sequon which displays the highest variability in occupancy and is most likely to directly influence the interaction with PD-1.

## Introduction

T cell mediated immune response is to a large extent controlled by co-stimulatory and co-inhibitory immune checkpoint signals (1). The programmed cell death receptor-1 (PD-1, *CD279*) and its ligand, programmed cell death ligand-1 (PD-L1, *CD274*), belong to the family of immune checkpoint inhibitors that under physiological circumstances maintain immune tolerance and regulate inflammatory responses to protect tissues from damage (2–4). Engagement of PD-1, expressed on a surface of antigen-activated T-lymphocytes (5), with its ligand PD-L1 ultimately leads to T cell exhaustion and immunosuppression (6, 7). This regulatory mechanism is frequently hijacked by cancer cells, usually through increased surface expression of PD-L1, which helps them evade immune surveillance (8, 9). For this reason, the two proteins became important targets of cancer immunotherapy (10).

Both PD-L1 and PD-1 are single pass transmembrane glycoproteins and recent publications provide evidence that the function of PD-L1 is regulated by its N-glycosylation (11, 12). N-glycosylation is a common modification occurring on NXS/T sequons of proteins and carried out by a complex enzymatic machinery in the endoplasmic reticulum and Golgi compartments (13). Glycosylation regulates various protein functions, including protein folding, intracellular trafficking as well as protein-protein interactions (14). PD-L1 carries four NXS/T sequons that can be N-glycosylated at N35, N192, N200 or N219 (11). It has been demonstrated that glycosylation stabilizes PD-L1 by creating a spatial hindrance (at N192, N200, and N219 positions) that prevents the interaction of PD-L1 with glycogen-synthase kinase 3β (GSK3β) and thus protects PD-L1 from phosphorylation-dependent proteasome degradation (11). In addition, PD-L1 glycosylation was shown to potentiate its interaction with PD-1 leading to suppressed vulnerability to T-cell toxicity in a triple negative breast cancer model (12). The latter study suggested that specific glycoforms, in particular those carrying poly-N-acetyllactosamine chains (polyLacNAc), may have substantial regulatory function in the PD-L1/PD-1 interaction (12). This is further supported by the observation that reducing PD-L1 glycosylation with 2-deoxyglucose or inhibition of polyLacNAc formation results in increased susceptibility of triple negative breast cancer cells to T-cell toxicity (12, 15). In addition, it was also suggested that binding of some antibodies to PD-L1 is altered by its N-glycosylation status (12, 16). The structure of PD-L1/PD-1 complex and PD-L1 and PD-1 complexes with therapeutic antibodies have been determined (17–19) but little is known about specific distribution of their glycoforms especially with regard to the potentially impactful polyLacNAc structures.

We have therefore optimized LC-MS/MS collision energies to resolve glycan structural motives on the PD-L1 glycoprotein (20, 21). Using this approach, we demonstrate that full-length PD-L1 produced in breast cancer cells contains high proportion of polyLacNAc N-glycans at the N219 sequon and that the cell surface glycoforms represent a limited subset of the whole pool of PD-L1 forms detectable in the cancer cells. We also demonstrate that glycosylation and glycosite occupancy of the full-length PD-L1 substantially differ from that of its truncated soluble variants previously used in PD-1/PD-L1 binding studies (12, 22–24) which calls for caution when selecting the proteins for functional studies.

## Experimental Section

### Reagents and antibodies

Human recombinant PD-L1 (Phe19-Thr239) with C-terminal 6xHis tag produced in HEK293 cells and human recombinant PD-L1-Fc chimera protein produced in mouse NS0 cells were obtained from R&D Systems, Minneapolis, MN (cat # 9049-B7 and 156-B7, respectively). Human PNGase F for N-deglycosylation was obtained from New England Biolabs, Ipswitch, MA (cat # B3704). Anti-human PD-L1 antibodies, Durvalumab (Imfinzi, Astra Zeneca), Avelumab (Bavencio, EMD Serona), and Atezolizumab (Tecentriq, Roche) were obtained from Oncology Pharmacy, Lombardi Comprehensive Cancer Center, Georgetown University, Washington, DC as discarded reagents from ongoing clinical studies. Donkey anti-human IgG-HRP was from Jackson ImmunoResearch (cat # 709-035-149).

### Cell cultures

Wild-type MDA-MB-231 cells (ATCC # HTB-26) were grown in DMEM-F12 (Corning) supplemented with 10% FBS (Gibco), and 50 μg/ml gentamycin (Sigma) at 37°C under humidified 5% CO_2_ atmosphere. MDA-MB-231 cell line stably expressing human PD-L1 was a kind gift from Dr. Men-Chie Hung, The University of Texas MD Anderson, Houston, TX (12). The culture conditions were as for wild-type cells but culture medium was supplemented with 10 μg/ml puromycin to maintain selection pressure.

### Transfection

The construct carrying the extracellular domain of PD-L1 (Phe19-Thr239) with C-terminal twin-Strep-10x His dual tag followed by a stop codon was synthesized using GeneArt Gene Synthesis service (Invitrogen, Carlsbad, CA) with *SgfI* and *MluI* restriction sites at the beginning and the end of the ORF sequence, respectively. For mammalian expression, the ORF was subcloned to pCM6-Entry vector (Origene, Rockville, MD) and transfected to wild-type MDA-MB-231 cells using Lipofectamine 3000 (Invitrogen) in serum-free OptiMEM medium (Invitrogen) according to manufacturer’s instructions. The conditioned media were harvested 72 hrs post-transfection and subjected to IMAC purification as described below.

### Immobilized metal affinity chromatography (IMAC)

Secreted form of the PD-L1 carrying C-terminal 10x His tag was purified using AKTA Start FPLC and Tricorn 5/100 column (both GE Healthcare) filled with HisPur Ni-NTA Resin (Thermo # 88223) at 1 ml/min flow. PBS supplemented with 300 mM NaCl was used as equilibration and binding buffer, washing (10 CV) was performed with 12.5 mM imidazole and elution (10 CV) with 250 mM imidazole; both in binding buffer. Eluted PD-L1 was concentrated and buffer exchanged into TBS, 0.01% Tween 20 using Amicon Centrifugal Filters with 10 kDa cut-off (EMD Millipore). Protein concentration was determined by a Micro BCA Assay (Thermo) and the purity was assessed to > 90% by SDS-PAGE as described below.

### Immunoaffinity purification of PD-L1

Secreted form of PD-L1 protein (Phe19-Thr239, R&D System # 9049-B7) was diluted in 50 mM Tris, pH 7.5, 100 mM NaCl, 1% Triton X-100, and 1 mM EDTA (TNTE) or spiked into cell lysates of wild-type MDA-MB-231 cells that were prepared as described below. MDA-MB-231 cells stably expressing full-length PD-L1 were scraped, pelleted at 500 × g for 10 min and lysed in TNTE buffer (1:10 wt:vol) supplemented with protease inhibitor cocktail (Roche) by homogenization using motorized Teflon-glass homogenizer (Thomas Scientific, 3000 rpm, 20 strokes). Lysates were incubated on ice for 30 min with frequent vortexing and cleared by centrifugation at 16,000 × g for 15 min at 4°C.

Durvalumab antibody was covalently immobilized to 4% beaded agarose using AminoLink Plus Immobilization Kit (Thermo Scientific # 44894) according to manufacturer’s instructions (4 mg antibody per 1 ml resin). The beads were equilibrated in TNTE buffer followed by overnight rotation at 4°C with PD-L1 protein or cleared lysate in ratio of 20 μl beads (dry volume)/ml lysate (buffer). The bead-sample suspension was transferred to Minispin columns (Thermo Fisher) and spun for 30 sec at 100 × g to collect flow-through. The beads were washed with 4 × 500 μl TBS, 0.1% Tween 20 followed by 4 × 500 μl wash with TBS. Bound PD-L1 was eluted with 100 μl 0.1 M glycine, pH 2.5 and immediately neutralized with 1 M Tris, pH 9. The eluate was either loaded on SDS-PAGE as described below or precipitated with acetone (1:5 vol:vol) and reconstituted in 50 mM ammonium bicarbonate buffer, pH 8 for trypsin digestion and LC-MS analysis as described below.

### Enrichment of membrane proteins by cell-surface biotinylation

MDA-MB-231 cells stably expressing full-length PD-L1 were grown in 175 cm^2^ flasks in complete DMEM-F12 medium as described above. Upon reaching 90% confluence the cells were incubated overnight in serum-free medium, washed twice with DPBS and incubated with 0.25 mg/ml of EZ-Link Sulfo-NHS-SS-Biotin cross-linking reagent (Thermo # 21331) in DPBS supplemented with 0.1 mM oxidized glutathione (Thermo Fisher) for 30 min at room temperature under constant rocking. The reaction was stopped by the addition of lysine to 50 mM final concentration. The cells were scraped, washed in TBS and resuspended in TNTE lysis buffer (see above) supplemented with a protease inhibitor cocktail (Roche). The suspension was homogenized by motorized Teflon-glass homogenizer (20 strokes, 3000 rpm), incubated on ice for 30 min with frequent vortexing and the homogenates were cleared by centrifugation at 16,000 g for 15 min at 4°C. Cleared homogenates were incubated overnight under constant rotation with High Capacity Neutravidin Beads (Thermo # 29202) equilibrated in lysis buffer. The beads were washed with TBS-T and bound biotinylated proteins were eluted by 60 min incubation with 50 mM dithiotreitol at 37°C. The individual fractions were analyzed by Western blot using Na+/K+ ATPase and PD-L1 antibodies as described below. The eluted samples were precipitated with acetone (1:5 vol:vol) and reconstituted in 50 mM bicarbonate buffer, pH 8 for trypsin digestion and LC-MS analysis as described below.

### SDS-PAGE and Western blot

Samples were resolved on 4-12% Bis-Tris NuPAGE gels (Invitrogen) according to manufacturer’s recommendations and either stained with Commasie Blue (Bulldog Bio, Portsmouth, NH) or transferred to PVDF membrane for Western Blot staining using iBlot 2 Gel Transfer Device (Invitrogen). Western blot was performed using iBind Automated Western System (Invitrogen) according to manufacturer’s instructions using human anti-PD-L1 antibody (Avelumab, EMD Serona, 1 μg/ml) or rabbit monoclonal Na/K ATPase antibody (Abcam # ab76020, 1:100,000) as primary antibodies and peroxidase-conjugated donkey anti-human IgG-HRP (Jackson ImmunoResearch, 1:2000) or goat anti-rabbit IgG-HRP (Abcam # ab6721, 1:10,000) as sencondary antibodies. Membranes were developed using Clarity Western ECL Substrate (Bio-Rad, Hercules, CA) and images were acquired using Amersham Imager 600 (GE Healthcare) with automated high dynamic range operation.

### Surface plasmon resonance (SPR)

N-deglycosylation of recombinant PD-L1 (R&D Systems) was performed using PNGase F (New England Biolabs) without a heat-inactivation step. Inactivated PNGase F (70°C for 10 min) was added to control samples. SPR was performed at 25°C using Biacore T200 system with CM5 chips (GE Healthcare). HBS-T (10 mM Hepes, pH 7.4, 150 mM NaCl, 0.005% Tween 20) was used as running buffer for all measurements. Surface of the CM5 sensor was pre-coated with 1 μM protein A/G using standard amine coupling chemistry to a level of approximately 4200 RU. Anti-PD-L1 antibodies, Avelumab, Durvalumab, and Atezolizumab, were captured onto the surface to a level of 50, 75, and 160 RU, respectively. The blank channel of the chip served as a negative control. Gradient concentrations of glycosylated (gPD-L1) and deglycosylated PD-L1 (dgPD-L1) (0 nM, 3.125 nM, 6.25 nM, 12.5 nM, 25 nM, and 50 nM) were flown over the chip surface. The flow rate of all analyte solutions was maintained at 50 μl/min. Association and dissociation times used for all analytes were 120 sec and 600 sec, respectively. All analytes were injected in triplicate. One 20 sec pulse of glycine, pH 1.5 was used to regenerate the surface of the sensor chip. Under these regeneration conditions, all ligands were dissociated from the protein A/G surface and fresh ligand solutions were injected every cycle before the analytes. The binding was analyzed by 1:1 kinetics model fitting.

### Glycopeptide preparation

Protein standards or isolated proteins after acetone precipitation were solubilized with 50 mM sodium bicarbonate as described above, reduced with 5 mM DTT for 60 min at 60°C and alkylated with 15 mM iodoacetamide for 30 min in the dark. Trypsin Gold (Promega, Madison, WI) digestion (2.5 ng/μl) was carried out at 37°C in Barocycler NEP2320 (Pressure BioSciences, South Easton, MA) for 1 hour.

### Glycopeptide analysis using DDA nano LC-MS/MS

Glycopeptides prepared as above were analyzed by liquid chromatography-tandem mass spectrometry (LC-MS/MS) using Orbitrap Fusion Lumos connected to a Dionex 3500 RSLC-nano-LC (Thermo Scientific) under workflow previously optimized for glycopeptide analysis (20). Briefly, separation of glycopeptides and peptides was achieved by a 5 min trapping/washing step using 99% solvent A (2% acetonitrile, 0.1% formic acid) at 10 μl/min followed by a 90 min acetonitrile gradient on a 150 mm × 75 μm C18 pepmap column at a flow rate of 0.3 μl/min: 0-3 min 2% B (0.1% formic acid in ACN), 3-5 min 2-10% B; 5-60 min 10-45% B; 60-65 min 45-98% B; 65-70 min 98% B, 70-90 min equilibration by 2% B. The electrospray ionization voltage was 3 kV and the capillary temperature 275°C. MS1 scans were performed over m/z 400–1800 with the wide quadrupole isolation on a resolution of 120,000 (m/z 200), RF Lens at 40%, intensity threshold for MS2 set to 2.0e4, selected precursors for MS2 were with charge state 3-8, and dynamic exclusion 30s. Data-dependent HCD tandem mass spectra were collected with a resolution of 15,000 in the Orbitrap with fixed first mass 110 and 5 normalized collision energy settings of 15, 20, 25, 30 and 35%.

### Glycopeptide identification

Byonic software (Protein Metric) was used for the identification of summary formulas of peptide-glycosylation. Independent searching was performed on the data with different collision energy settings as described recently (21). All spectra of identified glycopeptides were checked manually for the presence of structure-specific fragments and quantitative information was extracted using Qualbrowser (Thermo) and Peakview (Sciex) software. In addition, retention behavior of each occupied glycopeptide (glycosite) was confirmed by analysis of PNGase F deglycosylated sample under ^18^O water as we described (20). Site occupancy was determined from the sum of areas of identified glycopeptides and area of non-glycosylated peptide.

### Modeling of the structure of glycosylated PD-1/PD-L1 complex

The complex of human PD-1 and PD-L1 in PDBID 4ZQK (17) was used as the basis of 3D structures. The missing domain in hPD-L1 was generated by superimposing the full-length structure of hPD-L1 from PDBID 4Z18. The missing loop from hPD-1 (residues 54-61) was generated using Modeller as described (25). In-house code based on the GEMS/GMML molecular modeling C++ library was used to add the glycans M5GN2F1 or M5GN2 to each glycosylation sequon in hPD-L1 (Uniprot # Q9NZQ7, glycosylation sites N35, N192, N200, N219) and hPD-1 (Uniprot # Q15116, glycosylation sites N49, N58, N74, N116). All molecular dynamics (MD) simulations were performed using the Amber18 software (University of California, San Francisco, CA). Using the Amber software package tleap, the 3D structures were placed in a cubic box of TIP5P water with a 10 Å water buffer with counter ions added to neutralize the system. The FF14SB (26) and Glycam06k (27) force fields were employed for the protein and the carbohydrate, respectively. Non-bonded cut-offs of 10.0 Å for vdW and 8.0 Å for electrostatics were employed. Initial energy minimization (10,000 steps steepest decent followed by 10,000 steps conjugate gradient) was performed with Cartesian restraints (5 kcal/mol throughout all phases) on all solute heavy atoms to optimize the water molecules position and orientation. The same restraints were employed during 400 ps nPT equilibration phase at 300°K with a Langevin thermostat. This was followed by a 1 ns structural equilibration phase with Cartesian restraints on protein Cα atoms only. This structure was then used to start five independent (random seed) production phase simulations of 100 ns without Cartesian restraints. The Amber program cpptraj was used to calculate per-residue b-factors over the course of the simulations. The 3D structure images were generated using VMD (28) with the 3D-SNFG symbol script (29).

## Results

### LC-MS analysis of PD-L1 glycoforms

We have analyzed the glycoforms of PD-L1 using our recently introduced analytical workflow implementing collision energy (CE) modulation for the resolution of specific structural motifs of glycans (20, 21). We optimized collision energy for the analysis of each structural motif described. Our analyses show that, depending on the protein site and cell used for expression, the N-glycans are occupied by structures carrying LacdiNAc, polyLacNac, or Gal-Gal structural motifs in addition to the expected common complex glycoforms. Figure 1A shows low CE (NCE 20) fragmentation spectrum of the N35 glycopeptide of PD-L1 produced in HEK293 cells that carries a complex glycan with a fucosylated polyLacNAc structural motif. The presence of characteristic oxonium ions (m/z 731, 877, 893, 1062) confirms the polyLacNAc motif while fragments (m/z 512, 877) confirm outer arm fucosylation. The polyLacNAc motif of the N219 glycopeptide of PD-L1, produced in breast MDA-MB-231 cells, is documented in supplementary Figure S1 that compares low and high CE fragmentation spectra of the core fucosylated tetra-antennary complex glycan. The low collision (NCE 15) fragmentation yields polyLacNAc-specific ions (m/z 731, 1022) (Figure S1B) that are not visible in the spectrum obtained at high CE (NCE 35, Figure S1A). Figure 1B shows low collision energy (NCE15) HCD spectrum of the N35 glycopeptide carrying Galα1-3Gal (Galili) motif on the PD-L1 produced in mouse myeloma NS0 cells. The dominant fragment m/z 528 confirms the presence of the Galili structural motif while the peak m/z 674 confirms outer arm fucosylation. Figure 2 demonstrates the resolution of glycans carrying GalNAcβ1– 4GlcNAcβ1-R (LacdiNAc) motif on the N200 glycopeptide of PD-L1 produced in HEK293 cells (Figure 2A). Fragmentation of the LacdiNAc carrying glycopeptides produces a characteristic HexNAc-HexNAc (m/z 407) fragment with intensity comparable to the usual HexNAc-Hex (m/z 366) fragment (Figure 2A). This oxonium ion distinguishes LacdiNAc carrying glycopeptides from peptides carrying the isobaric multiantennary complex glycans. This is demonstrated in Figure 2B showing HCD fragmentation spectrum of the N200 glycopeptide of PD-L1 produced in breast cancer MDA-MB-231 cells where a dominant m/z 366 oxonium ion and lack of the m/z 407 fragment (Figure 2B) confirms a tetra-antennary agalactosylated structure. It is of note that the HexNAc-HexNAc oxonium ion can also be produced by fragmentation from the chitobiose core of any glycopeptide but the intensity of this fragment is negligible compared to the LacdiNAc containing glycans even at higher collision energies (NCE 35) (Figure 2A and B).

**Figure 1.**
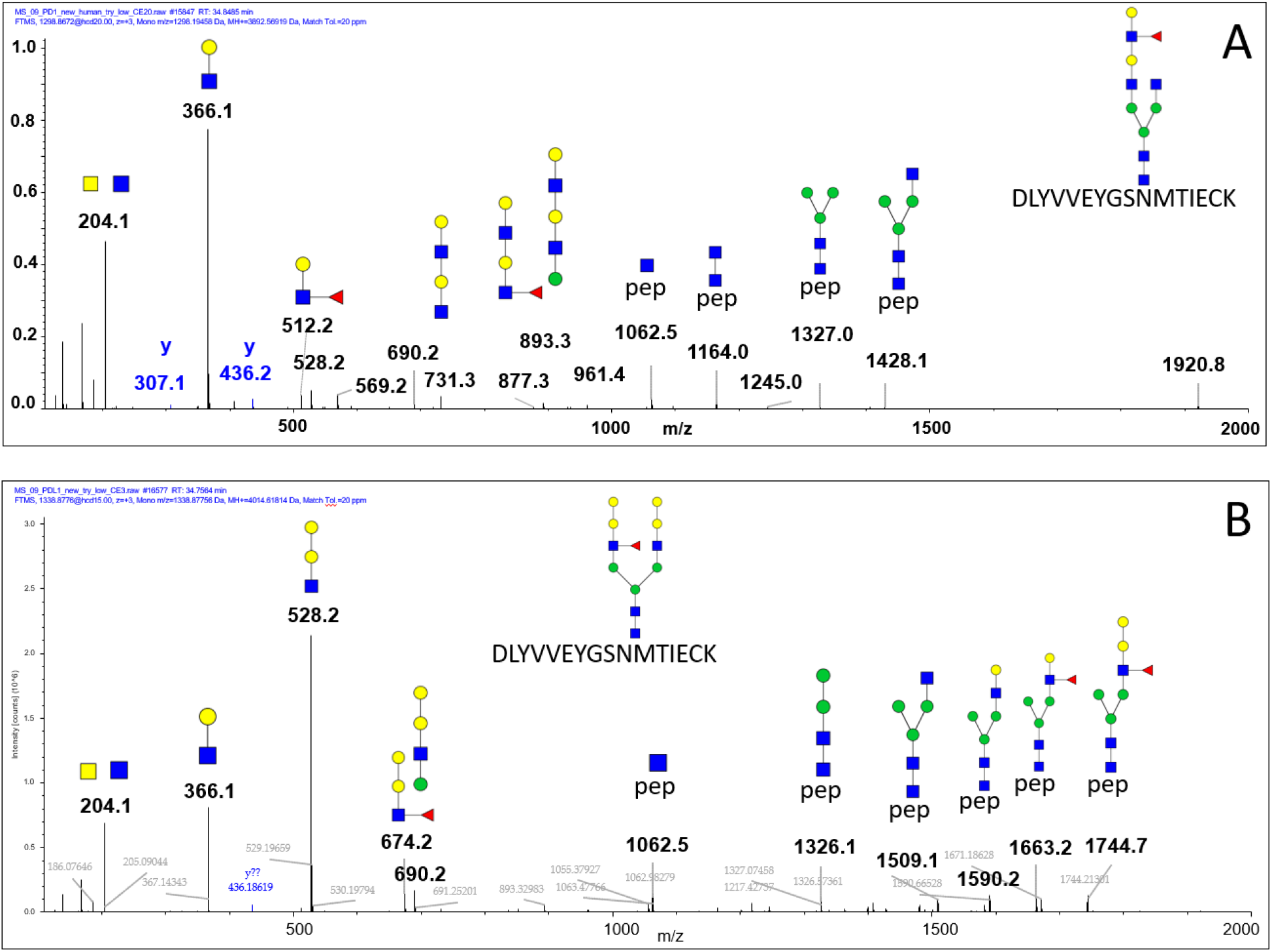
(**A**) Low collision (NCE 20) HCD tandem mass spectrum of PD-L1 glycopeptide N35 carrying the polyLacNAc motif with outer arm fucosylation and produced in HEK293 cells. (**B**) Low collision (NCE 15) spectrum of PD-L1 glycopeptide N35 containing Gal-Gal (Galili) motif and produced in mouse NS0 cells.

**Figure 2.**
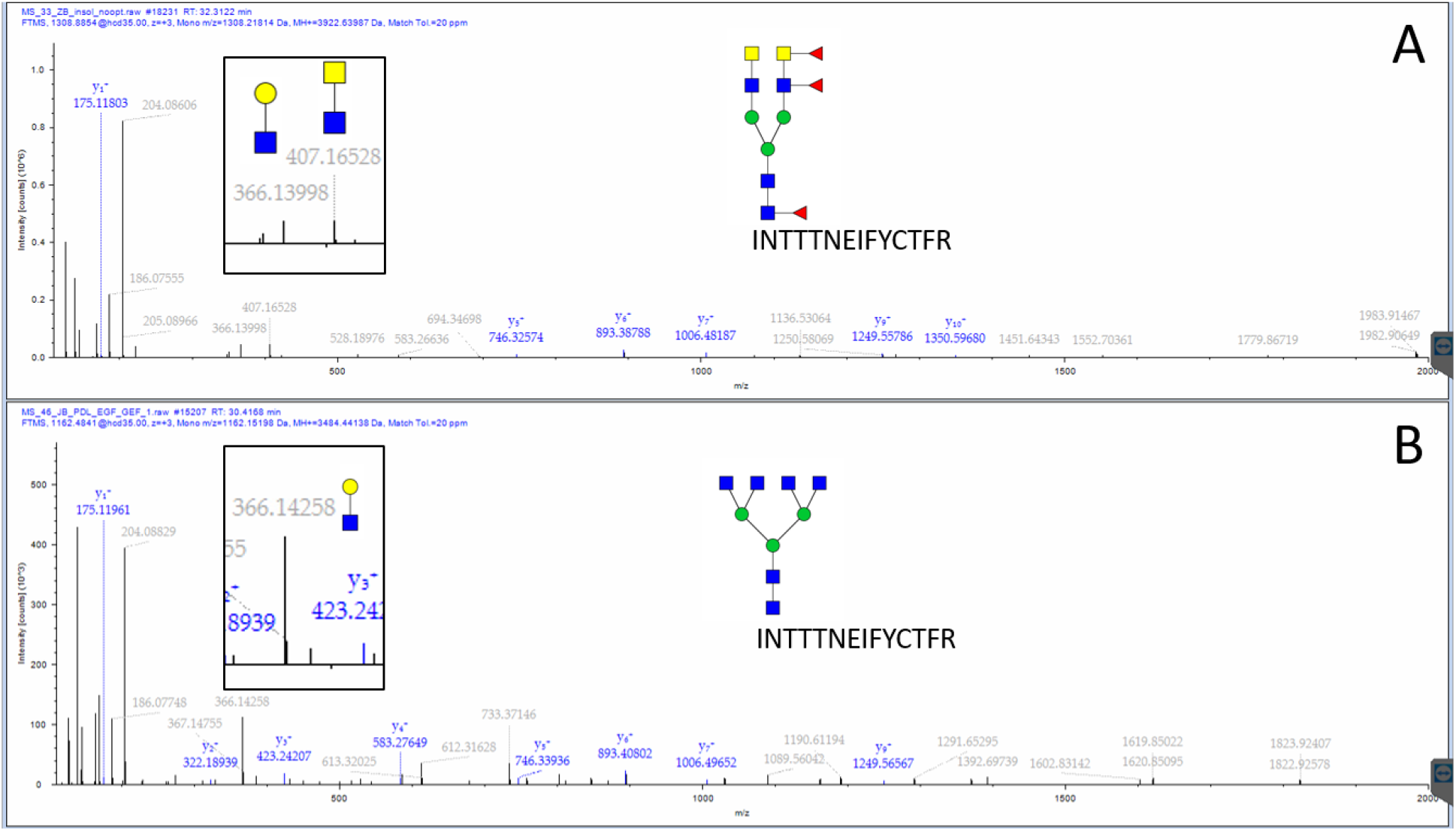
HCD tandem mass spectrum of PD-L1 glycopeptide N200 produced in HEK293 (**A**) and MDA-MB-231 (**B**) cells. HEK293-produced glycopeptide contains LacdiNAc structure motifs confirmed by the presence of the m/z 407 oxonium ion (insert A) as opposed to the N200 glycopeptide of PD-L1 produced in MDA-MB-231 cells where lack of m/z 407 fragment (insert B) confirms a tetra-antennary agalactosylated structure.

**Figure 3.**
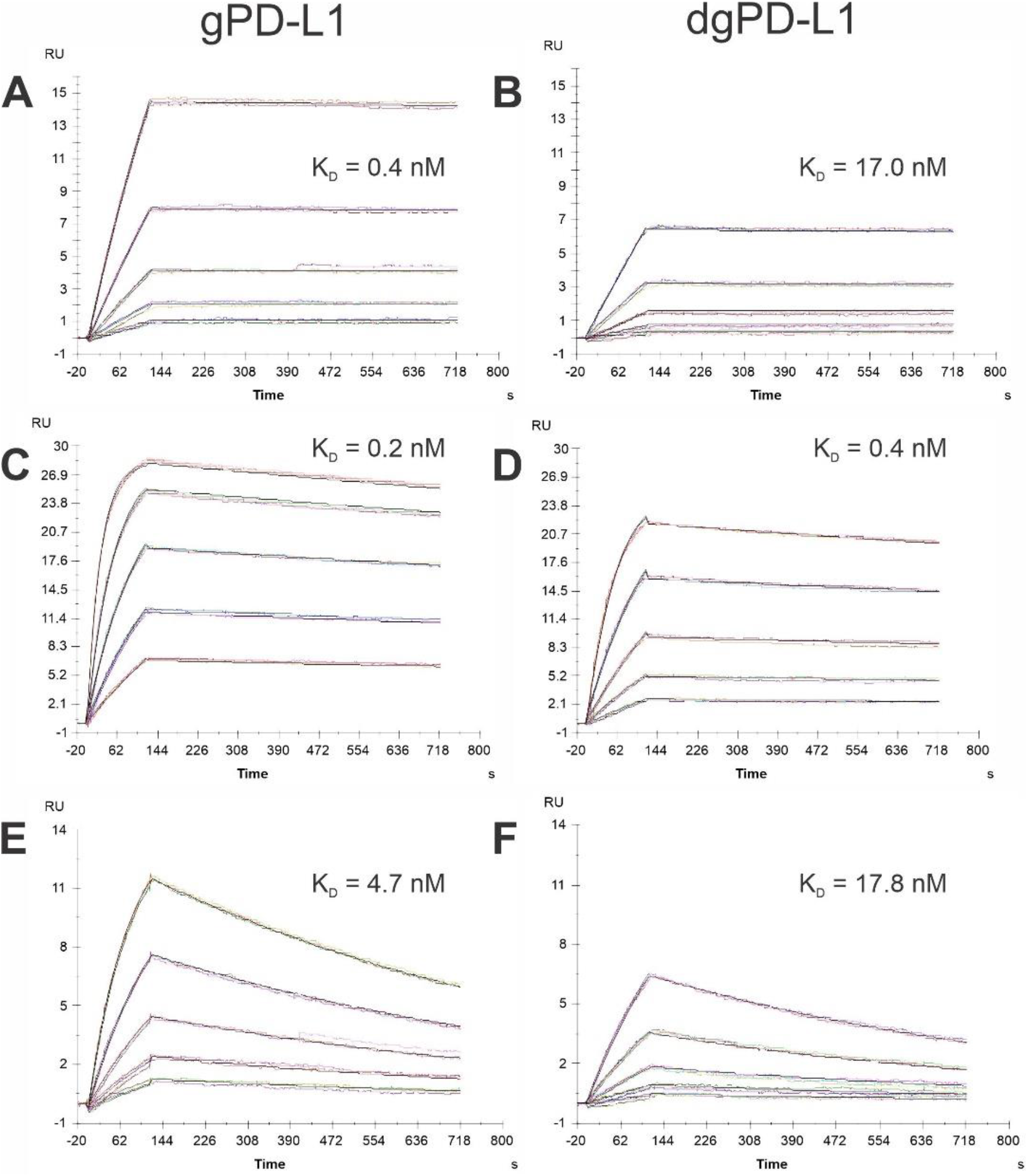
SPR characterization of the binding of glycosylated (gPD-L1) and deglycosylated PD-L1 (dgPD-L1) to the following antibodies: Avelumab (A, B), Durvalumab (C, D), and Atezolizumab (E, F). PD-L1 (analyte) concentrations used were 3.125 nM, 6.25 nM, 12.5 nM, 25 nM, and 50 nM.

### Comparison of the glycoforms of PD-L1 produced in human HEK293 and mouse NS0 cells

Two commercially available PD-L1 proteins lacking the transmembrane domain were utilized in previous studies aimed to determine the impact of PD-L1 glycosylation on interaction with PD-1 (12, 24). The truncated form of the protein (Phe19-Thr239) with C-terminal His tag was produced in human HEK293F cells (R&D Systems # 9049-B7) and the chimera protein with C-terminal Fc domain of human IgG1 (R&D Systems # 156-B7) was produced in mouse NS0 cells. The proteins were used to demonstrate that deglycosylation of PD-L1 substantially reduces its interaction with the PD-1 protein (12). We have analyzed the glycoforms of both proteins by energy optimized LC-MS/MS (21). We find that the two proteins substantially differ not only in glycoforms but also in site occupancy (Table 1). This is most clearly demonstrated at the N219 sequon which is almost fully occupied in PD-L1 produced in the mouse cells while less than 1% occupied in the PD-L1 expressed in the human HEK293 cells (Table 1). The N219 site had the highest proportion of complex glycans with polyLacNAc chains (Table 1, CP) previously suggested to be important for the interaction with PD-L1 *in vivo* (12). Other major difference was observed at the N192 sequon which was almost exclusively occupied by high mannose (HM) structures in the mouse cells in contrast to the HEK293-produced PD-L1. HEK293-produced PD-L1 had high proportion of complex glycans carrying the terminal GalNAcβ1-4GlcNAcβ1-R (LacdiNAc) motif (Table 1). On the other hand, PD-L1 from the mouse cells carried glycans containing the Gal-Gal (GG) motif (Table 1) not seen, as expected, in the glycoproteins from the human cells.

**Table 1.** Glycosylation of the secreted form of PD-L1 (Phe19-Thr239) produced by human HEK293, mouse NS0, and human MDA-MB-231 cells analyzed on four glycopeptides: DLYVVEYGSNMTIECK (N35), LFNVTSTLR (N192), INTTTNEIFYCTFR (N200), and LDPEENHTAELVIPELPLAHPPNER (N219). Abbreviations: HexNAc, N-acetylhexosamine; Hex, Hexose; Fuc, Fucose; SA, sialic acid; SO, site occupancy; GP#, number of identified glycopeptides; NQ, not quantifiable. Three most abundant glycoforms for each site are listed. Types of glycan structure: HM, high mannose; H, hybrid; C, complex; CL, complex glycan containing LacdiNAc; CP, complex glycan containing polyLacNAc; GG, complex glycan containing Gal-Gal (Galili); PM, paucimannose.

| Origin | Site | SO | GP# | Glycan | Type | % |
| --- | --- | --- | --- | --- | --- | --- |
| HEK293<br>(human) | N35 | >99% | 44 | HexNAc(6)Hex(4)SA(1) | CL | 13 |
|  |  |  |  | HexNAc(6)Hex(5) | CL | 8 |
|  |  |  |  | HexNAc(6)Hex(4)Fuc(1) | CL | 6 |
|  | N192 | >99% | 75 | HexNAc(4)Hex(4) | CL | 15 |
|  |  |  |  | HexNAc(4)Hex(4)Fuc(1) | CL | 13 |
|  |  |  |  | HexNAc(4)Hex(5) | CL | 12 |
|  | N200 | >81% | 61 | HexNAc(5)Hex(4)Fuc(1)SA(2) | CL | 19 |
|  |  |  |  | HexNAc(5)Hex(4)SA(2) | C | 18 |
|  |  |  |  | HexNAc(5)Hex(4)Fuc(2) | C | 5 |
|  | N219 | <0.1% | 0 | NQ | n/a | n/a |
| NS0<br>(mouse) | N35 | 98% | 7 | HexNAc(5)Hex(9)Fuc(1) | GG | 43 |
|  |  |  |  | HexNAc(5)Hex(8)Fuc(1) | GG | 30 |
|  |  |  |  | HexNAc(9)Hex(4) | CL | 14 |
|  | N192 | >99% | 19 | HexNAc(2)Hex(5) | HM | 47 |
|  |  |  |  | HexNAc(2)Hex(7) | HM | 32 |
|  |  |  |  | HexNAc(2)Hex(6) | HM | 17 |
|  | N200 | 97% | 17 | HexNAc(5)Hex(9)Fuc(1) | GG | 67 |
|  |  |  |  | HexNAc(4)Hex(7)Fuc(1) | GG | 5 |
|  |  |  |  | HexNAc(5)Hex(8)Fuc(1) | GG | 5 |
|  | N219 | >99% | 34 | HexNAc(8)Hex(8) | CP | 59 |
|  |  |  |  | HexNAc(8)Hex(7) | CP | 22 |
|  |  |  |  | HexNAc(8)Hex(6) | CP | 12 |
| MDA-MB-231<br>(human) | N35 | 91% | 24 | HexNAc(4)Hex(6)Fuc(1) | H | 26 |
|  |  |  |  | HexNAc(4)Hex(6)Fuc(2) | H | 23 |
|  |  |  |  | HexNAc(2)Hex(5) | HM | 10 |
|  | N192 | >99% | 34 | HexNAc(2)Hex(9) | HM | 25 |
|  |  |  |  | HexNAc(3)Hex(5)SA(1) | H | 16 |
|  |  |  |  | HexNAc(2)Hex(6) | HM | 12 |
|  | N200 | >99% | 30 | HexNAc(4)Hex(5)Fuc(1) | C | 43 |
|  |  |  |  | HexNAc(4)Hex(4)Fuc(1) | C | 12 |
|  |  |  |  | HexNAc(4)Hex(3)Fuc(1) | PM | 8 |
|  | N219 | <1% | 9 | HexNAc(5)Hex(5) | C | 33 |
|  |  |  |  | HexNAc(4)Hex(5)Fuc(3) | C | 18 |
|  |  |  |  | HexNAc(2)Hex(5) | HM | 17 |

### Glycoforms observed on the secreted form of PD-L1 (Phe19-Thr239) produced in the triple-negative breast cancer cell line MDA-MB-231

To compare the commercially available PD-L1 protein with the corresponding counterpart produced by breast cancer MDA-MB-231 cells, we transfected the cells with truncated PD-L1 (Phe19-Thr239) carrying C-terminal His tag and purified it from conditioned media to > 90% purity as described in the Experimental Section. We do this because the triple negative breast cancer cell lines MDA-MB-231 and the truncated recombinant PD-L1 proteins produced by human HEK-293 or mouse NS0 cells were used previously to demonstrate the impact of glycosylation on its binding with PD-1. Compared to the HEK-293 produced protein, PD-L1 from the breast cancer cells contained substantially higher proportion of high mannose (Table 1, HM) and hybrid (H) structures that represent majority of the glycoforms on the N35 and N192 sequons (Table 1). Another difference is the high proportion of complex glycans with LacdiNAc motif in PD-L1 from HEK293 cells that are not seen in breast cancer cell produced protein (Table 1, CL). On the other hand, the occupancy is similar in both cell lines with high occupancy at all sequons except N219 which is occupied less than 1% for both proteins. The above results suggest substantial differences in glycosylation of the truncated soluble form of PD-L1 (Phe19-Thr239) dependent on the cellular origin.

### Binding of PD-L1 antibodies to glycosylated and deglycosylated PD-L1

The extracellular domains of the membrane bound proteins are useful for the studies of protein-protein interactions *in vitro* but their glycoforms may not be representative of the natively occurring full-length protein incorporated in the membrane of relevant cells. For this reason, we wanted to compare the glycoforms of the truncated protein to the full-length PD-L1 expressed in the triple-negative breast cancer (TNBC) cells used in the studies of PD-L1 (11, 12, 15). To achieve this goal, we needed to enrich PD-L1 from TNBC cells and we used for this purpose high affinity therapeutic antibodies. Because affinity of the antibodies can be affected by the glycosylation of their targets (12, 16), we decided to test the influence of glycans on their binding to avoid selective enrichment of some PD-L1 glycoforms. We analyzed the interaction of glycosylated and deglycosylated PD-L1 with immobilized antibodies using surface plasmon resonance (SPR) assays (Figure 2). Of the three tested antibodies (Avelumab, Durvalumab, Atezolizumab), Durvalumab has shown the highest affinity toward PD-L1 (K_D_ 0.2 nM) and the lowest selectivity toward its glycosylated form (K_D_ 0.2 and 0.4 nM for glycosylated and deglycosylated PD-L1, respectively). In addition, as shown in Supplemental Table S1, association and dissociation rates for glycosylated and deglycosylated PD-L1 were comparable for the Durvalumab antibody (k*a* 73.7 vs 48.5 10^4^M^−1^s^−1^; k*d* 1.73 vs 1.86 10^−4^s^−1^; for glycosylated vs deglycosylated PD-L1, respectively) while differences were more pronounced for the remaining antibodies. The largest preference for glycosylated PD-L1 was observed for the Avelumab antibody (K_D_ 0.4 vs 17.0 nM, k*a* 6.74 vs 0.23 104M^−1^s^−1^, for glycosylated and deglycosylated PD-L1, respectively) (Supplemental Table 1). The lack of glycoform selection was further confirmed by comparison of the PD-L1 glycoforms before and after pull-down with the immobilized Durvalumab antibody (Supplemental Table S2). The results demonstrate that the distribution of glycoforms is not affected by the pull-down which further confirms suitability of Durvalumab for the glycosylation-independent enrichment of PD-L1.

### Glycosylation of full-length PD-L1 produced in triple negative breast cancer MDA-MB-231 cells

Since only cell surface PD-L1 protein participates in cell-cell interactions, we compared PD-L1 glycoforms from whole cell lysates and cell surface fraction of the MDA-MB-231 cells obtained by surface biotinylation as described in the Experimental Section. In the whole cell preparation (Total PD-L1) we observed significant proportion of high mannose glycans on all four glycopeptides (Table 2) with proportional contribution as follows: N35 (39%), N192 (100%), N200 (76%), and N219 (51%). This is in contrast with the cell surface PD-L1, where high mannose structures were observed exclusively at N192 (Table 2) which prefers high mannose glycans for unknown, perhaps structural, reasons. We identified considerable proportion of complex glycans with polyLacNAc structural epitope at the N219 sequon (Table 2, CP) that was substantially higher in the cell surface PD-L1 (37% vs 85%, respectively). PolyLacNAc glycans were previously proposed as an important determinant of the high affinity of glycosylated PD-L1 for PD-1 (12). High proportion of polyLacNAc structures and high occupancy of the N219 site (92% and 80% in total and surface PD-L1, respectively, Table 2) indicates that glycosylation at this sequon may indeed be regulated to adjust the interaction. High occupancy and large proportion of the polyLacNAc carrying glycans in the full-length membrane-bound protein is a major difference from its secreted form produced in the same cell line (Table 1). In addition, the cell surface protein carries substantially fewer glycoforms. The restricted presentation is not an artefact of analysis as we isolate sufficient quantities for the LC-MS/MS analysis by our efficient immunoaffinity procedure. The glycoforms presented at the cell surface are simply less heterogeneous than the glycoforms present in the total cell lysate and, especially, compared to the glycoforms on the secreted forms of the proteins. It appears that the membrane anchoring of the substrate limits the variability of the glycoforms produced and released to the cell surface.

**Table 2.** Glycosylation of the four glycopeptides of PD-L1 isolated from whole cell lysate and enriched cell-surface fraction of MDA-MB-231 cells. Abbreviations: HexNAc, N-acetylhexosamine; Hex, Hexose; Fuc, Fucose; SA, sialic acid; SO, site occupancy, GP#, number of identified glycopeptides. Types of glycan structure: HM, high mannose; C, complex; CP, complex glycan containing polyLacNAc

| Site | Total |  |  |  | Membrane |  |  |  |
| --- | --- | --- | --- | --- | --- | --- | --- | --- |
|  | SO<br>(GP#) | Glycan | Type | % | SO<br>(GP#) | Glycan | Type | % |
| **N35** | >99% | HexNAc(4)Hex(5)Fuc(1)SA(1) | C | 43 | >99% | HexNAc(4)Hex(5)Fuc(3) | C | 100 |
|  | (4) | HexNAc(2)Hex(9) | HM | 34 | (1) |  |  |  |
|  |  | HexNAc(4)Hex(5)Fuc(3) | C | 18 |  |  |  |  |
|  |  | HexNAc(2)Hex(9) | HM | 5 |  |  |  |  |
| **N192** | >99% | HexNAc(2)Hex(7) | HM | 46 | >99% | HexNAc(2)Hex(7) | HM | 47 |
|  | (6) | HexNAc(2)Hex(8) | HM | 25 | (5) | HexNAc(2)Hex(8) | HM | 42 |
|  |  | HexNAc(2)Hex(9) | HM | 14 |  | HexNAc(2)Hex(9) | HM | 8 |
|  |  | HexNAc(2)Hex(6) | HM | 7 |  | HexNAc(2)Hex(5) | HM | 2 |
|  |  | HexNAc(2)Hex(5) | HM | 2 |  | HexNAc(2)Hex(6) | HM | 1 |
|  |  | HexNAc(2)Hex(4) | HM | 1 |  |  |  |  |
| **N200** | >99% | HexNAc(2)Hex(9) | HM | 38 | >99% | HexNAc(4)Hex(5)Fuc(1)SA(2) | C | 40 |
|  | (6) | HexNAc(2)Hex(8) | HM | 32 | (4) | HexNAc(4)Hex(5)Fuc(1)SA(1) | C | 26 |
|  |  | HexNAc(4)Hex(5)Fuc(1)SA(1) | C | 12 |  | HexNAc(6)Hex(7)Fuc(1)SA(3) | C | 20 |
|  |  | HexNAc(4)Hex(5)Fuc(1)SA(2) | C | 6 |  | HexNAc(7)Hex(8)SA(1) | CP | 14 |
|  |  | HexNAc(2)Hex(7) | HM | 6 |  |  |  |  |
|  |  | HexNAc(5)Hex(6)Fuc(3)SA(1) | C | 4 |  |  |  |  |
| **N219** | 92% | HexNAc(2)Hex(9) | HM | 26 | 80% | HexNAc(5)Hex(6)Fuc(1)SA(2) | CP | 22 |
|  | (14) | HexNAc(2)Hex(8) | HM | 11 | (10) | HexNAc(7)Hex(8)SA(1) | CP | 19 |
|  |  | HexNAc(5)Hex(6)Fuc(1)SA(2) | CP | 8 |  | HexNAc(7)Hex(8)Fuc(1)SA(2) | CP | 18 |
|  |  | HexNAc(7)Hex(8)Fuc(1)SA(2) | CP | 8 |  | HexNAc(7)Hex(8)Fuc(1)SA(3) | CP | 16 |
|  |  | HexNAc(2)Hex(7) | HM | 7 |  | HexNAc(4)Hex(5)Fuc(1)SA(2) | C | 8 |
|  |  | HexNAc(5)Hex(6)Fuc(1)SA(1) | CP | 7 |  | HexNAc(5)Hex(6)Fuc(3) | CP | 5 |
|  |  | HexNAc(2)Hex(6) | HM | 6 |  | HexNAc(4)Hex(5)Fuc(1)SA(1) | C | 5 |
|  |  | HexNAc(6)Hex(7)Fuc(1)SA(2) | CP | 5 |  | HexNAc(5)Hex(6)Fuc(1)SA(1) | CP | 4 |
|  |  | HexNAc(7)Hex(8)SA(1) | CP | 4 |  | HexNAc(6)Hex(7)Fuc(1)SA(4) | C | 3 |
|  |  | HexNAc(4)Hex(5)Fuc(1)SA(2) | C | 3 |  | HexNAc(6)Hex(7)Fuc(1)SA(3) | CP | 1 |
|  |  | HexNAc(7)Hex(8)Fuc(1)SA(4) | CP | 3 |  |  |  |  |
|  |  | HexNAc(7)Hex(8)Fuc(1)SA(3) | CP | 2 |  |  |  |  |
|  |  | HexNAc(6)Hex(7)Fuc(1)SA(4) | C | 2 |  |  |  |  |
|  |  | HexNAc(2)Hex(5) | HM | 1 |  |  |  |  |

### Human PD-1/PD-L1 complex 3D structure modeling

The impact of glycosylation on PD-L1 stability was estimated from the average b-factors for the protein backbone atoms (Cα) for nude vs glycosylated PD-L1 with Man5 or Man5-Fuc glycans at each site. The average b-factors were as follows: 42.1 ± 10.7 for nude PD-L1, 35.1 ± 8.6 (p = 0.27, n = 5) for Man5-PD-L1, and 32.7 ± 5.3 for Man5Fuc-PD-L1 (p = 0.07, n = 5). These differences are not statistically significant, indicating that other factors than protein backbone dynamics of the PD-L1 fold may be involved in the effect of glycosylation on PD-L1 stability.

MD simulation of PD-1/PD-L1 complex with N-glycans is shown in Figure 4 for generic Man5-Fuc glycans (A). Overlaid images of 50 snapshots taken at regular intervals from the MD simulation show the extent of glycan motions (Figure 4B). The simulations of this complex showed transient interactions between the glycan at N219 of PD-L1 with those at N74 and N116 of PD-1. However, no discernable impact of glycans on the stability of the complex was observed. However, the glycan at N74 of PD-1 is located at the binding interface and the possibility that it stabilizes the complex cannot be eliminated solely on the basis of the MD data, which may only be manifest over much longer time scales than the present simulations.

**Figure 4.**
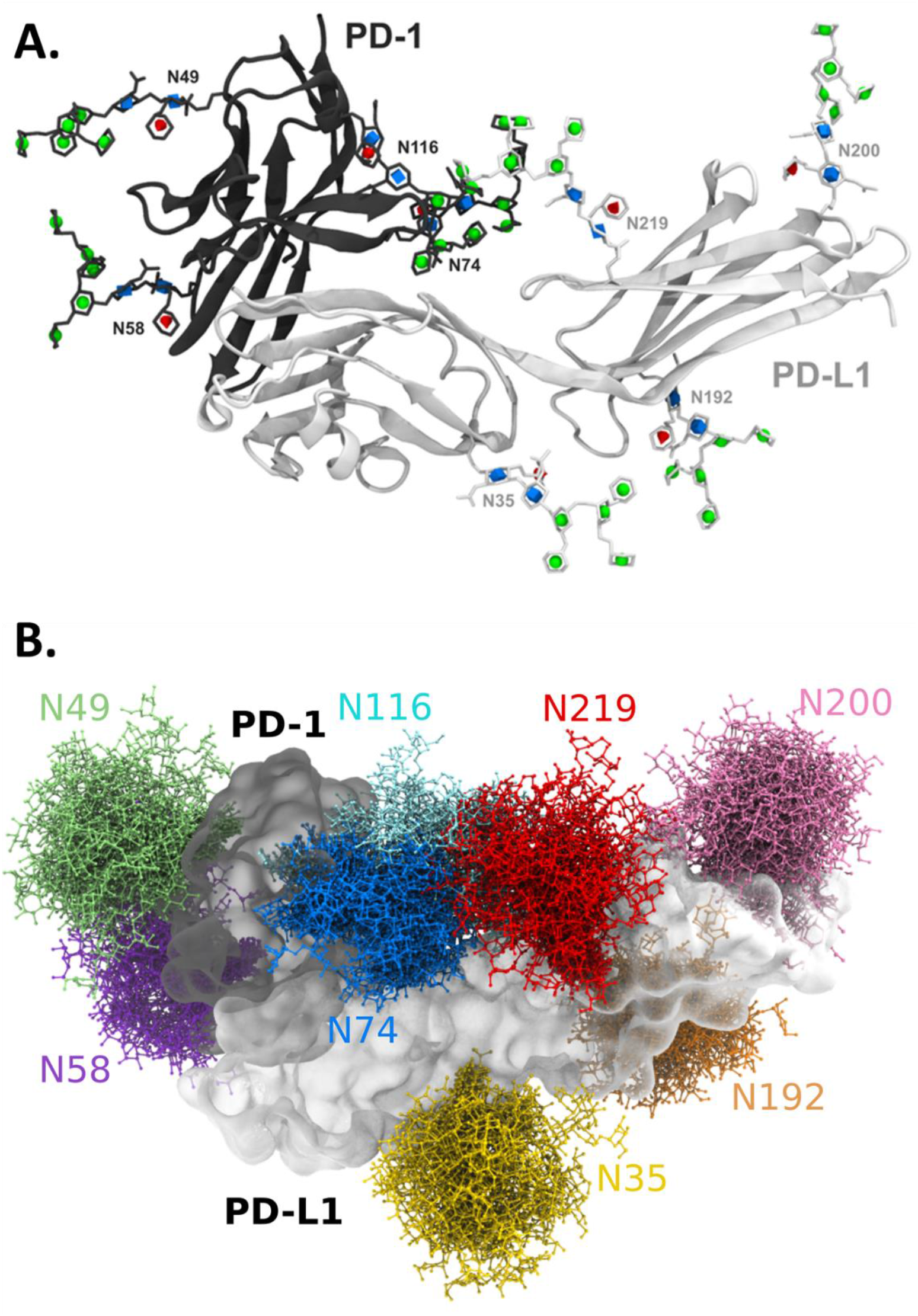
(**A**) Snapshot of the MD simulation of the glycosylated PD-L1/PD-1 complex using generic Man5-Fuc glycans. (**B**) Overlaid images from the MD showing the extent of glycan motions observed during the simulation. The models show that glycan N74 of PD-1 is proximal to the binding interface, and illustrates possible interactions between the N74 and N116 glycans on PD-1 with the N219 glycan of PD-L1.

## Discussion

N-glycosylation of PD-L1 has been recently suggested as a major regulator of the ability of breast cancer cells to avoid immune surveillance (11, 12). In particular, it was suggested that EGFR signaling drives expression of the beta-1,3-N-acetylglucosaminyltransferase 3 (*B3GNT3*) which leads to the formation of poly-N-acetyllactosamine (polyLacNAc) chains that enhance the interaction with PD-1 (12). Accordingly, knocking out *B3GNT3* in breast cancer cells reduced surface binding of PD-1 and sensitized cancer cells to T cell cytotoxicity (12) thus underlining the regulatory significance of the polyLacNAc carrying glycoforms. However, direct evidence of polyLacNAc-carrying glycans and their distribution across individual glycosylation sites was not reported. To provide detailed analysis of the glycoforms of PD-L1, we carried out optimized site-specific LC-MS/MS analysis of the PD-L1 protein expressed in several forms and in three different cell lines used in the previous studies.

We utilized immunoaffinity purification to achieve sufficient enrichment of PD-L1 from the cell lysates for our LC-MS/MS studies. We tested three clinically used antibodies for this purpose (Avelumab, Durvalumab, Atezolizumab) because of their documented strong binding and selectivity (30). However, the reported binding studies were performed with *E.coli* produced non-glycosylated PD-L1 and it was shown that affinity of some antibodies to PD-L1 is altered by its N-glycosylation status (12, 16). We therefore performed SPR analysis of the three clinical antibodies with glycosylated and de-glycosylated PD-L1 (Figure 2, Supplemental Table S1). Durvalumab demonstrated strongest binding and the lowest selectivity for glycosylated PD-L1 which was further confirmed by LC-MS analysis of glycoforms before and after pull-down (Supplemental Table S2). The remaining antibodies have shown certain preference for glycosylated PD-L1 that was most pronounced for Avelumab (K_D_ 0.4 vs 17.0 nM, k*a* 6.74 vs 0.23 10^4^M^−1^s^−1^, for glycosylated and deglycosylated PD-L1, respectively) (Supplemental Table S1). It is of note, however, that even though we observed forty-fold reduction in the affinity of Avelumab for the deglycosylated PD-L1, the affinity is still almost three orders of magnitude higher than the reported 8 μM K_D_ value for the PD-1/PD-L1 interaction (22, 23). Based on these results, it is not likely that glycosylation status of PD-L1 would significantly affect the therapeutic efficiency of these clinical antibodies. In addition, non-glycosylated/underglycosylated PD-L1 is unstable and is subject to GSK3β-mediated proteasome degradation (11) therefore it is unlikely that it will be exposed on the cell surface. However, analysis of PD-L1 glycoforms in the context of pathology, and in particular malignancies, may open new avenues for the development of glyco-specific antibodies that may reduce impact and risks of PD-L1 blockade in healthy tissues.

Analysis of complex glycans is complicated by the instability of the outer arm structural features under fragmentation conditions used in standard glycoproteomic workflows. We performed our analysis under LC-MS/MS conditions using limited collision energy fragmentation optimized to achieve resolution of some important structural motifs including the polyLacNAc containing N-glycopeptides (20, 21). Our results document the advantage of our optimized low CE fragmentation for structural resolution of the polyLacNAc glycoforms (Figure S1) and show that the polyLacNAc glycoforms represent a substantial component of the N-glycans at the N219 sequon. The limited CE fragmentation also enables resolution of other structural motifs of PD-L1 such as the outer-arm fucosylation (HexNAc-Hex-Fuc fragment m/z 512) or the LacdiNAc or Gal-Gal structural motives as described in detail previously (21).

We began our analysis with the truncated secreted form of the PD-L1(Phe19-Thr239). The truncated forms of membrane proteins are useful models to study protein function or protein-protein interactions *in vitro*. However, interaction between PD-L1 and PD-1 depends on their N-glycosylation and the question arises whether glycoforms of the recombinant model proteins, albeit produced in mammalian cells, reflect the membrane-integrated protein in the tumor cells with sufficient accuracy. Intracellular maturation of N-glycans into their mature forms is a complex enzymatic process carried out by numerous glycosyltransferases in the Golgi compartments and many factors, including spatial distribution of the enzymes and their substrates, determine which glycoforms are synthesized (31). Soluble proteins are spatially less restricted and may gain access to different glycosyltransferases than their membrane-bound counterparts. On the other hand, membrane integration of the glycosylated protein may be required to achieve conformation needed for access of a particular glycosyltransferase. Our results suggest that synthesis of the secreted PD-L1 indeed shifts the distribution of the glycoforms. The secreted PD-L1 expressed in the mouse NS0 or human HEK293 cells carries glycans typical of the cell of origin which is not surprising. However, all the secreted proteins, including protein expressed in the MDA-MB-231 cells, express wide variety of glycoforms compared to the full-length protein. Interestingly, while the total number of glycoforms is high, occupancy at some sites is lower in the secreted form, as seen for the N219 sequon in the human HEK293 and MDA-MB-231 cells, and some glycoforms are synthesized to a limited extent, for example the glycans carrying polyLacNAc chains. The secreted glycoproteins need to be therefore studied with caution.

The reason for the differences in N219 occupancy between full-length and truncated forms of the PD-L1 protein is not clear. The HEK293- and MDA-MB-231-produced truncated PD-L1 proteins are terminated shortly after Thr239 and carry only His tag sequence at their C-terminus. Oligosaccharyltransferase STT3A complex associated with the ER translocon mediates co-translational glycosylation on a nascent polypeptide chain entering the ER lumen (32, 33). It has been reported that NXS/T sequons within the last 65-75 residues of the polypeptide chain are more likely to be skipped by STT3A complex (34) thus explaining low occupancy at the N219 sequon in these proteins. This is further supported by the observation that PD-L1 chimera protein from mouse NS0 cells that is C-terminally extended by 238 aa IgG1-Fc domain shows high N219 occupancy (Table 1). Whether this is indeed the case or rather a cell type-specific feature needs to be determined by further studies. Such studies may be useful to reveal better approach to generate more relevant membrane protein models for *in vitro* studies.

Major focus of our study was to determine the glycoforms of PD-L1 bound to the plasma membrane of the triple negative breast cancer cell line MDA-MB-231. It was shown that these cells carry G13D KRas mutation (35) that leads to constitutive activation of the Ras-Raf-Mek-Erk signaling pathway, a major pathway of EGFR signaling in the breast cancer cells (36). Our results confirm that the MDA-MB-231 cells express the polyLacNAc glycoforms and that the expression is site-specific. It is primarily the N219 which carries this type of glycoforms and the polyLacNAc represents approximately 80% of the PD-L1 N-glycans at this site in the MDA-MB-231 membranes. Interestingly, polyLacNAc is rarely detected at other sequons and the N219 shows also the highest variability in occupancy. In combination, regulation of occupancy and polyLacNAc synthesis could be an important determinant of the PD-L1 interactions. The reason why polyLacNAc is preferentially formed at the N219 is not known but we know that B3GNT3 is a type II transmembrane protein (40). N219 of PD-L1 is the closest to the transmembrane region that begins at T239 of the PD-L1 sequence. It is possible that glycans at this sequon have the most favorable conformation for interaction with the B3GNT3 but further studies are needed to confirm this assumption.

The potential influence of the glycans at N219 on PD-L1 interactions is further supported by 3D modeling of the glycosylated PD-1/PD-L1 complex (Figure 4). The model suggests possible interactions between the N219 glycan of PD-L1 and the N74 and N116 glycans of PD-1. Based on the modeling results, PD-L1 folding is not significantly affected by glycosylation suggesting that direct interaction with N219 glycan might be the stabilizing feature of PD-1/PD-L1 interaction. However, further studies with appropriate protein models must be carried out to experimentally examine this possibility.

Cell surface PD-L1 is the form that actively regulates T-cell mediated immunosuppression and is therefore most relevant to analyze. We utilized surface biotinylation to capture the cell surface proteins and analysis of the surface fraction of PD-L1 revealed substantial differences of its glycoforms compared to the whole-cell lysate. Overall, the glycosylated sequons (N35, N192, N200 or N219) of the membrane integrated full-length PD-L1 isolated from the MDA-MB-231 triple negative breast cancer cells carry primarily complex N-glycans with the exception of N192 that is exclusively occupied by high mannose glycans (Table 2). This may be associated with lower accessibility of this sequon to the processing enzymes (37). PD-L1 from whole cell lysate contained high proportion of high mannose glycans (Table 2), generally considered to be an intermediate product of glycan maturation (38); the partially processed PD-L1 glycoforms likely reduce the relative contribution of polyLacNAc N-glycans in this preparation (Table 2). This observation suggests that substantial portion of cellular PD-L1 is stored intracellularly as immature glycoprotein. Given the potential importance of glycosylation in the regulation of PD-L1 function(s), it might be possible that intracellular immature PD-L1 serves as a pool that may be diverted into needed glycoforms in response to stimuli from the surrounding microenvironment.

## Conclusions

We observe substantial differences in glycosylation between the full-length native PD-L1 protein and its truncated extracellular variant (Phe19-Thr239) commonly used in *in vitro* studies. The most intriguing, and probably the most influential, differences are the low site occupancy at the N219 sequon and low content of glycans carrying polyLacNAc chains. Overall, we observe substantially higher variability of glycoforms of the truncated PD-L1. The membrane bound form of PD-L1 carries high proportion of polyLacNAc glycans at the N219 sequon which should be further evaluated as a means or regulation of its interactions in the immunological synapse. In addition, we demonstrate that, although clinically used antibodies differ in their preference for the glycosylated PD-L1, Durvalumab is an efficient tool for the isolation of PD-L1 from complex samples regardless of its glycoforms.

## Acknowledgements

We wish to thank Dr. Men-Chie Hung, The University of Texas MD Anderson, Houston, TX for kind gift of MDA-MB-231 cell line stably expressing human PD-L1. We thank Dr. Purushottam Tiwari and Prof. Aykut Uren from Genomics and Epigenomics Shared Resource, Lombardi Comprehensive Cancer Center, Georgetown University, Washington, DC for SPR analysis. Research reported in this publication was supported by the National Institutes of Health under awards S10OD023557, U01 CA230692, R01CA135069, and R01CA238455 to RG and CCSG Grant P30 CA51008. R.J.W. thanks the National Institutes of Health (U01 CA207824 and P41 GM103390) for financial support. The content is solely the responsibility of the authors and does not necessarily represent the official views of the National Institutes of Health.

## REFERENCES

1. Pardoll, D. M. (2012) The blockade of immune checkpoints in cancer immunotherapy. Nat. Rev. Cancer. 12, 252–264

2. Ceeraz, S., Nowak, E. C., and Noelle, R. J. (2013) B7 family checkpoint regulators in immune regulation and disease. Trends Immunol. 10.1016/j.it.2013.07.003

3. Pentcheva-Hoang, T., Corse, E., and Allison, J. P. (2009) Negative regulators of T-cell activation: potential targets for therapeutic intervention in cancer, autoimmune disease, and persistent infections. Immunol. Rev. 229, 67–87

4. Jung, K., and Choi, I. (2013) Emerging Co-signaling Networks in T Cell Immune Regulation. Immune Netw. 13, 184–193

5. Ishida, Y., Agata, Y., Shibahara, K., and Honjo, T. (1992) Induced expression of PD-1, a novel member of the immunoglobulin gene superfamily, upon programmed cell death. EMBO J. 11, 3887–3895

6. Freeman, G. J., Long, A. J., Iwai, Y., Bourque, K., Chernova, T., Nishimura, H., Fitz, L. J., Malenkovich, N., Okazaki, T., Byrne, M. C., Horton, H. F., Fouser, L., Carter, L., Ling, V., Bowman, M. R., Carreno, B. M., Collins, M., Wood, C. R., and Honjo, T. (2000) Engagement of the PD-1 immunoinhibitory receptor by a novel B7 family member leads to negative regulation of lymphocyte activation. J. Exp. Med. 192, 1027–1034

7. Topalian, S. L., Drake, C. G., and Pardoll, D. M. (2015) Immune checkpoint blockade: a common denominator approach to cancer therapy. Cancer Cell. 27, 450–461

8. Curiel, T. J., Wei, S., Dong, H., Alvarez, X., Cheng, P., Mottram, P., Krzysiek, R., Knutson, K. L., Daniel, B., Zimmermann, M. C., David, O., Burow, M., Gordon, A., Dhurandhar, N., Myers, L., Berggren, R., Hemminki, A., Alvarez, R. D., Emilie, D., Curiel, D. T., Chen, L., and Zou, W. (2003) Blockade of B7-H1 improves myeloid dendritic cell-mediated antitumor immunity. Nat. Med. 9, 562–567

9. Dong, H., Strome, S. E., Salomao, D. R., Tamura, H., Hirano, F., Flies, D. B., Roche, P. C., Lu, J., Zhu, G., Tamada, K., Lennon, V. A., Celis, E., and Chen, L. (2002) Tumor-associated B7-H1 promotes T-cell apoptosis: a potential mechanism of immune evasion. Nat. Med. 8, 793–800

10. Gong, J., Chehrazi-Raffle, A., Reddi, S., and Salgia, R. (2018) Development of PD-1 and PD-L1 inhibitors as a form of cancer immunotherapy: a comprehensive review of registration trials and future considerations. J Immunother Cancer. 6, 8

11. Li, C.-W., Lim, S.-O., Xia, W., Lee, H.-H., Chan, L.-C., Kuo, C.-W., Khoo, K.-H., Chang, S.-S., Cha, J.-H., Kim, T., Hsu, J. L., Wu, Y., Hsu, J.-M., Yamaguchi, H., Ding, Q., Wang, Y., Yao, J., Lee, C.-C., Wu, H.-J., Sahin, A. A., Allison, J. P., Yu, D., Hortobagyi, G. N., and Hung, M.-C. (2016) Glycosylation and stabilization of programmed death ligand-1 suppresses T-cell activity. Nat Commun. 7, 12632

12. Li, C.-W., Lim, S.-O., Chung, E. M., Kim, Y.-S., Park, A. H., Yao, J., Cha, J.-H., Xia, W., Chan, L.-C., Kim, T., Chang, S.-S., Lee, H.-H., Chou, C.-K., Liu, Y.-L., Yeh, H.-C., Perillo, E. P., Dunn, A. K., Kuo, C.-W., Khoo, K.-H., Hsu, J. L., Wu, Y., Hsu, J.-M., Yamaguchi, H., Huang, T.-H., Sahin, A. A., Hortobagyi, G. N., Yoo, S. S., and Hung, M.-C. (2018) Eradication of Triple-Negative Breast Cancer Cells by Targeting Glycosylated PD-L1. Cancer Cell. 33, 187–201.e10

13. Stanley, P., Taniguchi, N., and Aebi, M. (2015) N-Glycans. in Essentials of Glycobiology, 3rd Ed. (Varki, A., Cummings, R. D., Esko, J. D., Stanley, P., Hart, G. W., Aebi, M., Darvill, A. G., Kinoshita, T., Packer, N. H., Prestegard, J. H., Schnaar, R. L., and Seeberger, P. H. eds), Cold Spring Harbor Laboratory Press, Cold Spring Harbor (NY), [online] http://www.ncbi.nlm.nih.gov/books/NBK453020/ (Accessed May 4, 2020)

14. Schwarz, F., and Aebi, M. (2011) Mechanisms and principles of N-linked protein glycosylation. Curr. Opin. Struct. Biol. 21, 576–582

15. Shao, B., Li, C.-W., Lim, S.-O., Sun, L., Lai, Y.-J., Hou, J., Liu, C., Chang, C.-W., Qiu, Y., Hsu, J.-M., Chan, L.-C., Zha, Z., Li, H., and Hung, M.-C. (2018) Deglycosylation of PD-L1 by 2-deoxyglucose reverses PARP inhibitor-induced immunosuppression in triple-negative breast cancer. Am J Cancer Res. 8, 1837–1846

16. Lee, H.-H., Wang, Y.-N., Xia, W., Chen, C.-H., Rau, K.-M., Ye, L., Wei, Y., Chou, C.-K., Wang, S.-C., Yan, M., Tu, C.-Y., Hsia, T.-C., Chiang, S.-F., Chao, K. S.C., Wistuba, I. I., Hsu, J. L., Hortobagyi, G. N., and Hung, M.-C. (2019) Removal of N-Linked Glycosylation Enhances PD-L1 Detection and Predicts Anti-PD-1/PD-L1 Therapeutic Efficacy. Cancer Cell. 36, 168–178.e4

17. Zak, K. M., Kitel, R., Przetocka, S., Golik, P., Guzik, K., Musielak, B., Dömling, A., Dubin, G., and Holak, T. A. (2015) Structure of the Complex of Human Programmed Death 1, PD-1, and Its Ligand PD-L1. Structure. 23, 2341–2348

18. Lee, J. Y., Lee, H. T., Shin, W., Chae, J., Choi, J., Kim, S. H., Lim, H., Won Heo, T., Park, K. Y., Lee, Y. J., Ryu, S. E., Son, J. Y., Lee, J. U., and Heo, Y.-S. (2016) Structural basis of checkpoint blockade by monoclonal antibodies in cancer immunotherapy. Nat Commun. 7, 13354

19. Tan, S., Zhang, H., Chai, Y., Song, H., Tong, Z., Wang, Q., Qi, J., Wong, G., Zhu, X., Liu, W. J., Gao, S., Wang, Z., Shi, Y., Yang, F., Gao, G. F., and Yan, J. (2017) An unexpected N-terminal loop in PD-1 dominates binding by nivolumab. Nat Commun. 8, 14369

20. Benicky, J., Sanda, M., Brnakova Kennedy, Z., and Goldman, R. (2019) N-Glycosylation is required for secretion of the precursor to brain-derived neurotrophic factor (proBDNF) carrying sulfated LacdiNAc structures. J. Biol. Chem. 294, 16816–16830

21. Sanda, M., Benicky, J., and Goldman, R. (2020) Low collision energy fragmentation in structure-specific glycoproteomics analysis. Anal. Chem. 10.1021/acs.analchem.0c00519

22. Cheng, X., Veverka, V., Radhakrishnan, A., Waters, L. C., Muskett, F. W., Morgan, S. H., Huo, J., Yu, C., Evans, E. J., Leslie, A. J., Griffiths, M., Stubberfield, C., Griffin, R., Henry, A. J., Jansson, A., Ladbury, J. E., Ikemizu, S., Carr, M. D., and Davis, S. J. (2013) Structure and Interactions of the Human Programmed Cell Death 1 Receptor. J. Biol. Chem. 288, 11771–11785

23. Lin, D. Y., Tanaka, Y., Iwasaki, M., Gittis, A. G., Su, H.-P., Mikami, B., Okazaki, T., Honjo, T., Minato, N., and Garboczi, D. N. (2008) The PD-1/PD-L1 complex resembles the antigen-binding Fv domains of antibodies and T cell receptors. PNAS. 105, 3011–3016

24. Sun, L., Li, C.-W., Chung, E. M., Yang, R., Kim, Y.-S., Park, A. H., Lai, Y.-J., Yang, Y., Wang, Y.-H., Liu, J., Qiu, Y., Khoo, K.-H., Yao, J., Hsu, J. L., Cha, J.-H., Chan, L.-C., Hsu, J.-M., Lee, H.-H., Yoo S. S., and Hung, M.-C. (2020) Targeting glycosylated PD-1 induces potent anti-tumor immunity. Cancer Res. 10.1158/0008-5472.CAN-19-3133

25. Eswar, N., Webb, B., Marti-Renom, M. A., Madhusudhan, M. S., Eramian, D., Shen, M., Pieper, U., and Sali, A. (2006) Comparative Protein Structure Modeling Using Modeller. Curr Protoc Bioinformatics. 0 5, Unit-5.6

26. Maier, J. A., Martinez, C., Kasavajhala, K., Wickstrom, L., Hauser, K. E., and Simmerling, C. (2015) ff14SB: Improving the Accuracy of Protein Side Chain and Backbone Parameters from ff99SB. J Chem Theory Comput. 11, 3696–3713

27. Kirschner, K. N., Yongye, A. B., Tschampel, S. M., González-Outeiriño, J., Daniels, C. R., Foley, B. L., and Woods, R. J. (2008) GLYCAM06: a generalizable biomolecular force field. Carbohydrates. J Comput Chem. 29, 622–655

28. Humphrey, W., Dalke, A., and Schulten, K. (1996) VMD: visual molecular dynamics. J Mol Graph. 14, 33-38, 27–28

29. Thieker, D. F., Hadden, J. A., Schulten, K., and Woods, R. J. (2016) 3D implementation of the symbol nomenclature for graphical representation of glycans. Glycobiology. 26, 786–787

30. Tan, S., Liu, K., Chai, Y., Zhang, C. W.-H., Gao, S., Gao, G. F., and Qi, J. (2018) Distinct PD-L1 binding characteristics of therapeutic monoclonal antibody durvalumab. Protein Cell. 9, 135–139

31. Thaysen-Andersen, M., and Packer, N. H. (2012) Site-specific glycoproteomics confirms that protein structure dictates formation of N-glycan type, core fucosylation and branching. Glycobiology. 22, 1440–1452

32. Nilsson, I., Kelleher, D. J., Miao, Y., Shao, Y., Kreibich, G., Gilmore, R., von Heijne, G., and Johnson, A. E. (2003) Photocross-linking of nascent chains to the STT3 subunit of the oligosaccharyltransferase complex. J. Cell Biol. 161, 715–725

33. Ruiz-Canada, C., Kelleher, D. J., and Gilmore, R. (2009) Cotranslational and posttranslational N-glycosylation of polypeptides by distinct mammalian OST isoforms. Cell. 136, 272–283

34. Shrimal, S., Trueman, S. F., and Gilmore, R. (2013) Extreme C-terminal sites are posttranslocationally glycosylated by the STT3B isoform of the OST. J. Cell Biol. 201, 81–95

35. Hollestelle, A., Elstrodt, F., Nagel, J. H.A., Kallemeijn, W. W., and Schutte, M. (2007) Phosphatidylinositol-3-OH kinase or RAS pathway mutations in human breast cancer cell lines. Mol. Cancer Res. 5, 195–201

36. Kim, R.-K., Suh, Y., Yoo, K.-C., Cui, Y.-H., Kim, H., Kim, M.-J., Gyu Kim, I., and Lee, S.-J. (2015) Activation of KRAS promotes the mesenchymal features of basal-type breast cancer. Exp Mol Med. 47, e137

37. Hang, I., Lin, C., Grant, O. C., Fleurkens, S., Villiger, T. K., Soos, M., Morbidelli, M., Woods, R. J., Gauss, R., and Aebi, M. (2015) Analysis of site-specific N-glycan remodeling in the endoplasmic reticulum and the Golgi. Glycobiology. 25, 1335–1349

38. Moremen, K. W., Tiemeyer, M., and Nairn, A. V. (2012) Vertebrate protein glycosylation: diversity, synthesis and function. Nat. Rev. Mol. Cell Biol. 13, 448–462

