## Supplemental Information for "PD-L1 glycosylation and its impact on binding to clinical antibodies"

| Ligand | Analyte | $K_D$ (nM) | $k_a$ ($10^4$/Ms) | $k_d$ ($10^{-4}$/s) |
|---|---|---|---|---|
| Avelumab | gPD-L1 | 0.4 | 6.74 | 0.29 |
| | dgPD-L1 | 17.0 | 0.23 | 0.39 |
| Durvalumab | gPD-L1 | 0.2 | 73.73 | 1.73 |
| | dgPD-L1 | 0.4 | 48.53 | 1.86 |
| Atezolizumab | gPD-L1 | 4.7 | 23.57 | 10.98 |
| | dgPD-L1 | 17.8 | 6.87 | 12.25 |

**Supplemental Table S1.** Kinetic parameters of interaction between antibodies and glycosylated (gPD-L1) or deglycosylated PD-L1 (dgPD-L1) analyzed by SPR assay with immobilized PD-L1 antibodies. $K_D$ – dissociation constant, $k_a$ – association rate constant, $k_d$ – dissociation rate constant.

| | | Before | | | After | | |
|---|---|---|---|---|---|---|---|
| Site | 3 major structures | SO | GP# | % | SO | GP# | % |
| **N35** | HexNAc(6)Hex(4)SA(1) | >99% | 44 | 13 | >99% | 73 | 14 |
| | HexNAc(6)Hex(5) | | | 8 | | | 5 |
| | HexNAc(6)Hex(4)Fuc(1) | | | 6 | | | 8 |
| **N192** | HexNAc(4)Hex(4) | 100% | 75 | 15 | >99% | 60 | 13 |
| | HexNAc(4)Hex(4)Fuc(1) | | | 13 | | | 12 |
| | HexNAc(4)Hex(5) | | | 12 | | | 11 |
| **N200** | HexNAc(5)Hex(4)Fuc(1)SA(2) | 81% | 61 | 19 | 76% | 38 | 20 |
| | HexNAc(5)Hex(4)SA(2) | | | 18 | | | 18 |
| | HexNAc(5)Hex(4)Fuc(2) | | | 5 | | | 2 |
| **N219** | --- | <0.1% | NQ | NQ | <0.1% | NQ | NQ |
| | --- | | | | | | |
| | --- | | | | | | |

**Supplemental Table S2.** Glycosylation of the secreted form of PD-L1 (Phe19-Thr239) produced in HEK293 cells and analyzed on four glycopeptides before and after immunoaffinity purification. The results show top 3 structures identified on PD-L1 before/after purification. Abbreviations: HexNAc, N-acetylhexosamine; Hex, Hexose; Fuc, Fucose; SA, sialic acid; GP#,.number of identified glycopeptides; NQ, not quantifiable; SO, site occupancy; GP#, number of identified glycopeptides

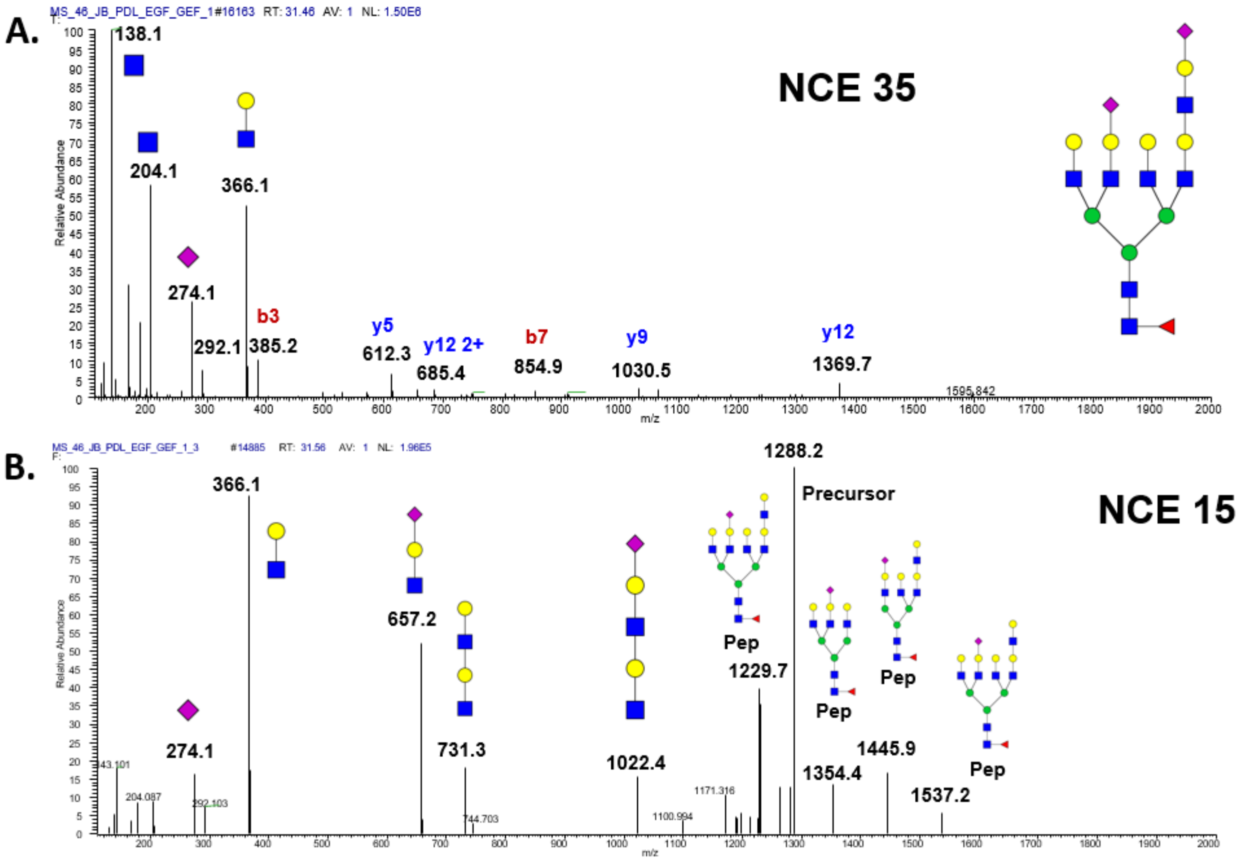


**Figure S1.** High (NCE 35) and low collision (NCE15) HCD tandem mass spectra of PD-L1 glycopeptide N219 produced in MDA-MB-231 cells. Series of peptide backbone fragment ions confirm occupied glycopeptide N219 in high collision energy spectra (**A**) and polyLacNAc-specific oxonium ions confirm the presence of polyLacNAc outer arm structure motif (**B**).
